# Succinate accumulation links mitochondrial MnSOD depletion to aberrant nuclear DNA methylation and altered cell fate

**DOI:** 10.1101/2020.06.24.169342

**Authors:** Kimberly L. Cramer-Morales, Collin D. Heer, Kranti A. Mapuskar, Frederick E. Domann

## Abstract

Previous studies showed that human cell line HEK293 lacking mitochondrial superoxide dismutase (MnSOD) exhibited decreased succinate dehydrogenase (SDH) activity, and mice lacking MnSOD displayed significant reductions in SDH and aconitase activities. Since MnSOD has significant effects on SDH activity, and succinate is a key regulator of TET enzymes needed for proper differentiation, we hypothesized that *SOD2* loss would lead to succinate accumulation, inhibition of TET activity, and impaired erythroid precursor differentiation. To test this hypothesis, we genetically disrupted the *SOD2* gene using the CRISPR/Cas9 genetic strategy in a human erythroleukemia cell line (HEL 92.1.7) capable of induced differentiation toward an erythroid phenotype. Cells obtained in this manner displayed significant inhibition of SDH activity and ~10-fold increases in cellular succinate levels compared to their parent cell controls. Furthermore, *SOD2*^−/−^ cells exhibited significantly reduced TET enzyme activity concomitant with decreases in genomic 5-hmC and corresponding increases in 5-mC. Finally, when stimulated with δ-aminolevulonic acid (δ-ALA), *SOD2−/−* HEL cells failed to properly differentiate toward an erythroid phenotype, likely due to failure to complete the necessary global DNA demethylation program required for erythroid maturation. Together, our findings support the model of an SDH/succinate/TET axis and a role for succinate as a retrograde signaling molecule of mitochondrial origin that significantly perturbs nuclear epigenetic reprogramming and introduce MnSOD as a governor of the SDH/succinate/TET axis.

## Introduction

Epigenetic processes provide a mechanism for integrating metabolic inputs and translating them into various phenotypic outputs [1,2]. Indeed, epigenetic modifiers often require or are inhibited by key citric acid cycle (CAC) metabolites (*e.g*. succinate, fumarate, alpha-ketoglutarate, acetyl-CoA) and co-factors (*e.g.* NAD+, FAD) [3]. One such metabolite, succinate, has recently been shown to act as a mitochondrial retrograde signal that exerts effects on the epigenome *via* inhibition of histone demethylases and the ten-eleven translocase (TET) family of DNA demethylase enzymes [4,5].

TET family enzymes contribute to the maintenance of the epigenome by catalyzing the oxidation of 5-methylcytosine (5-mC) to 5-hydroxymethylcytosine (5-hmC), initiating a series of oxidation reactions necessary for the removal of 5-mC. Like other epigenetic modifiers, TET family enzymes are sensitive to metabolic perturbations affecting the absolute and relative concentrations of their co-factors and products. Their required cofactors and substrates are identical to those of the collagen and HIF1 proline hydroxylase (PHD) family of enzymes and include molecular oxygen (O_2_), ferrous iron (Fe^2+^), α-ketoglutarate (α-KG), and ascorbate [6]. Using an FeIV-oxo complex generated by the oxidative decarboxylation of α-KG, TET enzymes catalyze the abstraction of a hydrogen from 5-mC, which is then hydroxylated. In the process, the oxidative decarboxylation of α-KG yields succinate and carbon dioxide (CO_2_) [7]. Through a product-induced negative-feedback mechanism, the inhibition of TET enzymes by succinate allows these enzymes to respond directly to changes in cellular metabolism, particularly to nodes linked to succinate and α-KG production and consumption.

Succinate and α-KG serve as key regulators of self-renewal and differentiation through TET enzymes. However, this regulation is complex and context-dependent [8,9]. For example, naïve embryonic stem cells sustain a high α-KG: succinate ratio to facilitate DNA and histone demethylation required for the maintenance of pluripotency [10,11]. However, α-KG is also important for the facilitation of epigenetic remodeling during differentiation of primed pluripotent stem cells [12] and premalignant cells in which p53 has been activated [13]. In the case of erythroid precursors, the model used in this study, global demethylation is required for the differentiation into red blood cells (RBCs) [14] and loss of TET enzymatic activity results in promoter hypermethylation and impaired differentiation of embryonic and hematopoietic stem cells [12,15,16]. Thus, the metabolic ratio of succinate to α-KG is a key factor in determining self-renewal and differentiation through modulation of TET activity.

A primary regulator of succinate levels is the activity of succinate dehydrogenase (SDH). SDH is a 4-subunit complex located in the inner membrane of the mitochondria that plays a role in the CAC *via* catalysis of the oxidation of succinate to fumarate as well as serving the role of Complex II in the electron transport chain by passing electrons from succinate or FADH_2_ directly to coenzyme Q in the mitochondrial inner membrane. Pharmacological inhibition or genetic mutation of SDH leads to accumulation of succinate and inhibition of α-KG-dependent enzymes such as TETs [5,17–19]. SDH activity can also be modulated by reactive oxygen species (ROS). Specifically, superoxide (O_2_^•-^) can inactivate iron-sulfur (Fe-S) cluster-containing enzymes such as SDH [18,19], as well as other enzymes in the CAC (aconitase, fumarate hydratase) [20,22,23], heme synthesis (ferrochelatase; FECH)[24], and amino acid metabolism [25,26] among several others.

Manganese superoxide dismutase (MnSOD) catalyzes the dismutation of mitochondrial O_2_^•-^ to hydrogen peroxide (H_2_O_2_). Cells lacking MnSOD exhibit elevated steady state levels of O_2_^•-^ and reduced SDH activity, likely due to inactivation of Fe-S clusters [20,27–29]. Since succinate is a direct regulator TET enzymatic activity and MnSOD has significant effects on SDH activity, we hypothesized that MnSOD loss would lead to succinate accumulation, inhibition of TET enzyme activity and altered cellular phenotype. To test this hypothesis, we used the CRISPR/Cas9 strategy to genetically disrupt the *SOD2* gene in two different human cell lines, human embryonic kidney 293 (HEK293) generated in a previous study [28], and a human erythroleukemia (HEL) generated in this study, thereby eliminating MnSOD activity. We chose HEL cells as a model because RBC maturation to produce mature enucleated erythrocytes requires rapid and widespread demethylation of genomic DNA [14] and these cells can be induced to differentiate toward mature red cells upon supplementation with δ-aminolevulinic acid (δ-ALA). Thus, we could utilize this model to study the effects of MnSOD loss on DNA demethylation-dependent erythroid differentiation.

Following characterization of the genetically disrupted *SOD2* cell lines, we measured the activities of mitochondrial enzymes sensitive to inactivation by O_2_^•-^(SDH, aconitase), succinate levels, TET enzyme activity, TET substrate (5-mC) and product (5-hmC) levels, and ability of HEL cells to undergo erythroid maturation in response to δ-ALA exposure. We found that MnSOD loss resulted in decreased SDH activity, succinate accumulation, reduced activity of TET enzymes, inhibition of DNA demethylation, and impaired erythroid differentiation. Taken together, our results introduce a mechanism through which MnSOD exerts epigenetic control over differentiation through an MnSOD/SDH/succinate/TET axis, and further support a role for succinate as a retrograde signaling molecule of mitochondrial origin.

## Materials and Methods

### Cell Lines

HEK293T cells were obtained from American Type Culture Collection (ATCC) maintained in DMEM supplemented with 10% fetal bovine serum in a humidified 37°C incubator with 5% CO_2_ and were routinely sub-cultured before reaching confluence by detachment with TrypLE Express (Invitrogen, Carlsbad, CA).

HEL 92.1.7 cells were obtained from ATCC and cultured in RPMI1640 supplemented with 10% fetal bovine serum in a humidified 37°C incubator with 5% CO_2_ and were routinely sub-cultured by 1:10 dilutions in fresh media.

### Plasmid Construction and Transfection

The pD130-GFP expression vector was designed to contain expression cassettes for green fluorescent protein (GFP), Cas9 endonuclease, and a CRISPR chimeric cDNA with the gRNA moiety designed to target exon 3 of SOD2 [ATATCAATCATAGCATTTTC]. This plasmid was custom synthesized (DNA2.0, Menlo Park, CA) and following amplification, purification, and characterization was used to transfect the two cell lines described above.

HEK293T cells were transiently transfected by calcium phosphate precipitation. Five days after transfection GFP+ cells were sorted and collected by flow cytometry and colonies were selected from single clones using cloning discs.

HEL 92.1.7 cells were transfected by electroporation at various conditions using the Gene Pulser Xcell™ Electroporation System (BioRad, USA). In brief, 5 × 10^6^ cells were suspended in 500 μl complete RPMI medium, mixed with 30 μg of pD130-GFP-plasmid in a 4 mm electroporation cuvette, and incubated on ice for 10 min. After electroporation with a single pulse (electrical parameters of 300 V, 950 μF) the cells were transferred into complete RPMI medium at a density of 1 × 10^6^ cells per ml and returned to the incubator. Five days after transfection GFP+ cells were sorted by flow cytometry and colonies were selected from single clones seeded in 96-well plates.

### Western Blot Analyses

Western blots were performed as previously described [26]. Briefly, total proteins were extracted from HEL 92.1.7 clones with standard RIPA buffer plus protease inhibitors and quantified by Bradford assay. Proteins were separated by SDS-PAGE, transferred to nitrocellulose membranes and probed with primary antibodies detecting MnSOD (Cat # 06-984, Millipore, Billerica, MA) and beta-tubulin (University of Iowa Hybridoma Core, Iowa City, IA).

### Aconitase activity

Aconitase activity was measured as described previously [30]. In brief, in a spectrophotometric assay where total cellular protein was combined with NADP^+^ and isocitrate dehydrogenase and the appearance of NADPH was monitored at 340 nm every 30s for 5 minutes.

### Succinate Dehydrogenase Histochemistry

Succinate dehydrogenase activity visualized via NBT reduction histochemistry. Culture medium was aspirated and cells were washed twice with PBS. Cell culture dishes were allowed to air dry at room temperature. A solution containing 0.55 mM NBT and 0.05 M sodium succinate was applied and the dishes were incubated overnight at 37°C and washed again with PBS before images were captured.

### Determination of Succinate levels

Succinate levels were determined using a colorimetric assay (Sigma-Aldrich Cat # MAK184) according to the manufacturer’s specifications.

### Tet DNA Demethylase Activity and 5-Hydroxy-Methylcytosine (5hmC) Measurement

Cellular TET DNA demethylase activities were assessed using the EpiGentek TET activity assay platform. Total genomic 5-methylcytosine and 5-hydroxymethylcytosine levels were measured in genomic DNA with the corresponding immunoassay systems also from EpiGentek according to the manufacturer’s instructions.

### Flow Cytometry Analysis

Determinations were performed with a flow cytometer (FACScan, Becton Dickinson, San Jose, CA). HEK293T or HEL 92.1.7 cells were gated on the basis of their forward scatter and side scatter signals with logarithmic scales. The region representing single, non-aggregated cells was gated before each experience with non-agglutinated samples to count events in the single-cell region. Fluorescence profiles were collected only on those cells appearing in this region to exclude agglutinates and debris. For each sample, 20,000 events were collected in logarithmic scale. Photomultiplier tube voltage for the fluorescence detector was adjusted before running samples to obtain a negative fluorescence signal for control (untransfected) samples. This source power value was fixed for all measurements. Computer software (WinMDI, version 2.8, Windows multiple document interface, flow cytometry application, Microsoft, Redmond, WA) was used for data analysis to calculate mean fluorescence intensity (MFI) from fluorescence histograms.

Antibodies utilized were directed against TFRC (PE-tagged anti-CD71 clone M-A712, BD Biosciences, San Jose, CA) and Glycophorin A (PE-tagged-anti-CD235a clone REA175, Miltenyi Biotech, Auburn, CA), markers of human erythroid maturation. Cells were incubated with antibodies under conditions previously described [31–33] and flow cytometry was performed as described above.

## Results

### Characterization of MnSOD loss in CRISPR/Cas9-edited HEL 92.1.7 cells

The CRISPR/Cas9 system for creating specific genetic deletions was used to target exon 3 of human *SOD2* in a HEL 92.1.7 cells. *SOD2*^−/−^ HEK293T were generated previously [28]. Two clones from each cell line were confirmed to show absence of MnSOD protein expression by Western blot analysis (**Figure 1**) [28]. In a previous study, we showed that MnSOD-deficient 293T (*SOD2*^−/−^) cells showed significantly increased DHE and MitoSOX fluorescence compared to wild-type (WT) cells, suggesting increased cytosolic and mitochondrial pro-oxidants levels, respectively [28].

**Figure 1.**
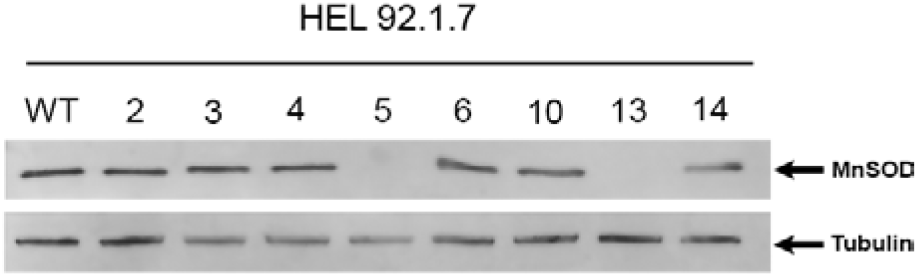
Characterization of MnSOD loss in CRISPR/Cas9-edited HEL 92.1.7 cells.

### *SOD2 KO cells exhibited decreased activity in enzymes sensitive to inactivation by* O_2_^•-^*and accumulation of cellular succinate*

Having established and characterized suitable model systems, we next sought to determine if *SOD2*^−/−^ cells exhibited decreased activity in mitochondrial enzymes known to be sensitive to inactivation by O_2_^•-^. Aconitase and SDH/Complex II are key players in mitochondrial metabolism that contain Fe-S clusters susceptible to inactivation by O_2_^•-^[20–22]. Aconitase activity was measured in a spectrophotometric assay where total cellular protein was combined with NADP^+^ and isocitrate dehydrogenase and the appearance of NADPH was measured at 340 nm. As expected, we observed a significant decrease in aconitase activity in *SOD2*^−/−^ HEL cells, likely due to inactivation by increased steady state levels of O_2_^•-^(**Figure 2A**). In addition, SDH activity was significantly decreased in *SOD2*^−/−^ HEL cells (**Figure 2B**) as was also observed in *SOD2*^−/−^ HEK293 cells in a previous study [28]. Consistent with the observed decrease in SDH activity, cellular succinate levels increased ~5-fold in both 293T and HEL KO cells (**Figure 2C**). Taken together, these results indicate that MnSOD loss is associated with significantly decreased SDH activity, likely through O_2_^•-^-mediated disruption of Fe-S clusters in SDH subunit B, and that loss of SDH activity is co-incident with increased levels of cellular succinate, the substrate for SDH.

**Figure 2.**
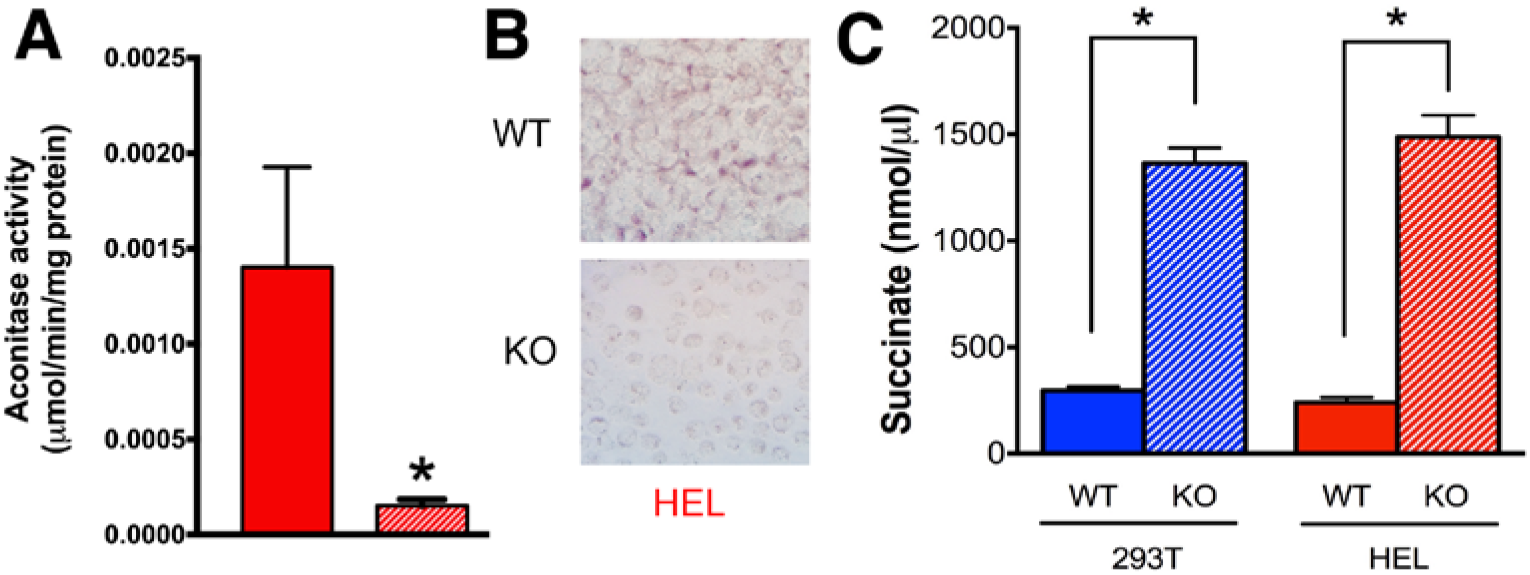
SOD2 KO cells exhibit decreased activity in enzymes sensitive to inactivation by superoxide and an accumulation of cellular succinate.

### Activity of TET family of demethylation enzymes was diminished in cells without MnSOD expression

TET DNA demethylases are nuclear enzymes that utilize α-KG and generate succinate, by which they known to be product inhibited. Therefore, we next sought to determine if the accumulation of succinate that occurs in *SOD2*^−/−^ cells coincided with decreased activity of TET family enzymes. To this end, TET activity was measured in an antibody-mediated colorimetric assay using nuclear protein. The results of this assay revealed a ~2-fold reduction in TET family activity in both *SOD2*^−/−^ 293T cells (blue striped bar) and SOD2−/− HEL cells (red striped bar) (**Figure 3**) compared to their WT parent cell types. Overall, these results demonstrate a clear association between loss of MnSOD expression and decreased activity of TET demethylase enzymes, likely due to product-induced negative feedback inhibition resulting from elevated succinate levels.

**Figure 3.**
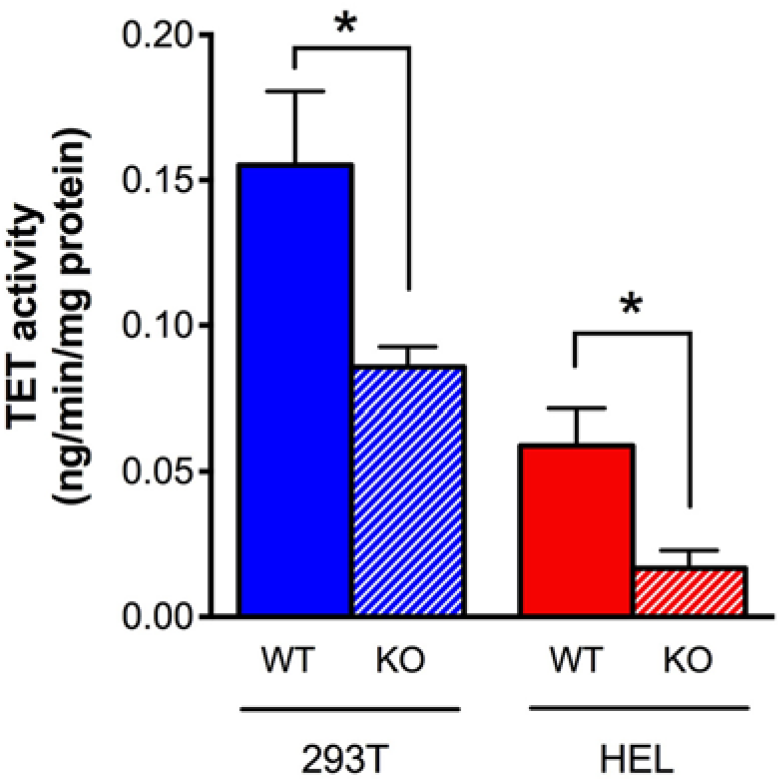
Activity of TET family of demethylation enzymes is diminished in cells without MnSOD expression.

### TET enzyme inactivation disrupted epigenetic programming and altered cell fate

Based on the observation of decreased TET enzyme activity in *SOD2*^−/−^ cell lines, we hypothesized that this would constitute a DNA demethylation defect and that these cells would therefore display an increase in the percentage of genomic 5-mC. Indeed, we observed a statistically significant increase in percent 5-mC in both 293T and HEL *SOD2*^−/−^ cells compared to their respective WT parent cell lines (**Figure 4A**). In parallel, we observed a corresponding decrease in the percentage of the TET enzyme product 5-hmC in HEL cells lacking MnSOD expression (**Figure 4B**). These results agree well with the finding of decreased TET enzyme activity and demonstrate that decreased TET enzyme activity is associated with an increase in TET enzyme substrate (5-mC) and decrease in TET enzyme product (5-hmC) potentially contributing to dysregulated homeostasis of DNA methylation

**Figure 4.**
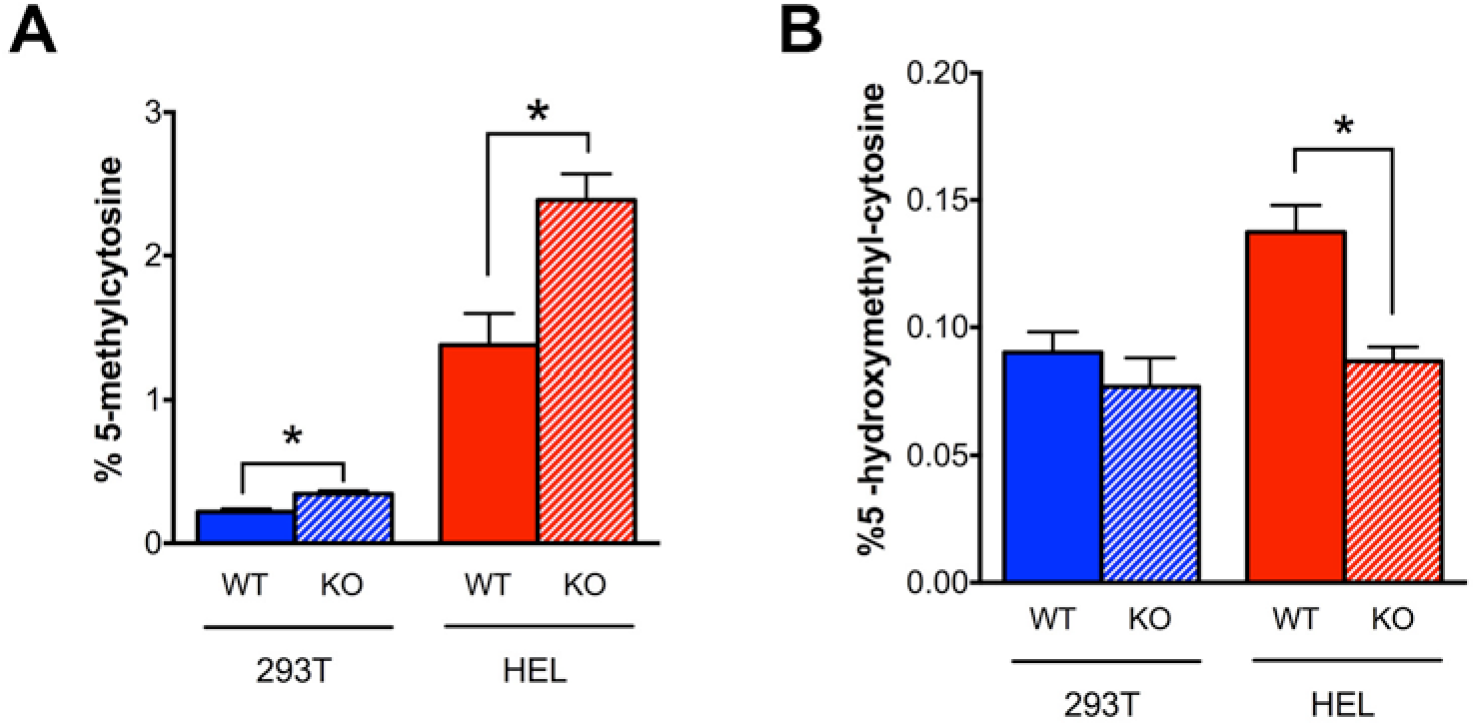
TET enzyme inactivation subsequently perturbs global epigenetic programming.

The process of erythroid maturation is dependent on global DNA demethylation, which we demonstrated was disrupted in *SOD2*^−/−^ cells. To determine whether the DNA demethylation defect in *SOD2*^−/−^ cells would impair the ability of erythroid-like precursor cells to differentiate toward RBCs, we compared the ability of parent WT HEL cells to *SOD2*^−/−^ HEL cells to differentiate upon exposure to δ-aminolevulinic acid (δ-ALA). Traditional markers (CD235a/Glycophorin A and CD71/Transferrin receptor) were used to assess erythroid maturation of HEL cells by flow cytometry. Differentiating RBC precursors display a wave of elevated CD71 expression that diminishes upon complete maturation to RBCs, and mature human erythroid cells are CD235a-positive and CD71-negative [34,35]. At baseline, *SOD2*^−/−^ cells exhibited slightly higher CD71 and CD235a expression compared to WT cells.

To initiate erythroid maturation, HEL cells were grown in the presence or absence of 100μM δ-aminolevulinic acid (δ-ALA) for 5 days. After incubation with δ-ALA for 5 days, WT HEL cells exhibited relatively low CD71 surface expression, consistent with a mature CD71-negative phenotype (**Figure 5, top**). By contrast, HEL *SOD2*^−/−^ cells (clone 5, blue and clone 13, orange) exhibited 70-80 fold increases in CD71 surface expression upon treatment with ALA demonstrating a significant defect in their ability to downregulate CD71 upon completion of the RBC maturation process (**Figure 5, top**). In the case of Glycophorin/CD235a, parent WT HEL cells upregulated CD235a 4-fold upon exposure to δ-ALA, consistent with normal erythroid maturation (**Figure 5, bottom**). In contrast, *SOD2*^−/−^ HEL cells exhibited a 2-4 fold decrease in CD235a (**Figure 5 bottom**). These results are quantified in **Table 1**. After 5 days in ALA, parent WT HEL cells resemble maturing erythroid cells (low TFRC, increased GlyA) whereas *SOD2*^−/−^ cells displayed aberrant expression of these biomarkers (high TFRC, decreased GlyA), consistent with their defect in DNA demethylation which is required for proper erythroid differentiation. Thus, *SOD2*^−/−^ HEL cells fail to display proper erythroid maturation. We propose that this impairment is likely due to a deficiency in TET-driven DNA demethylation.

**Figure 5.**
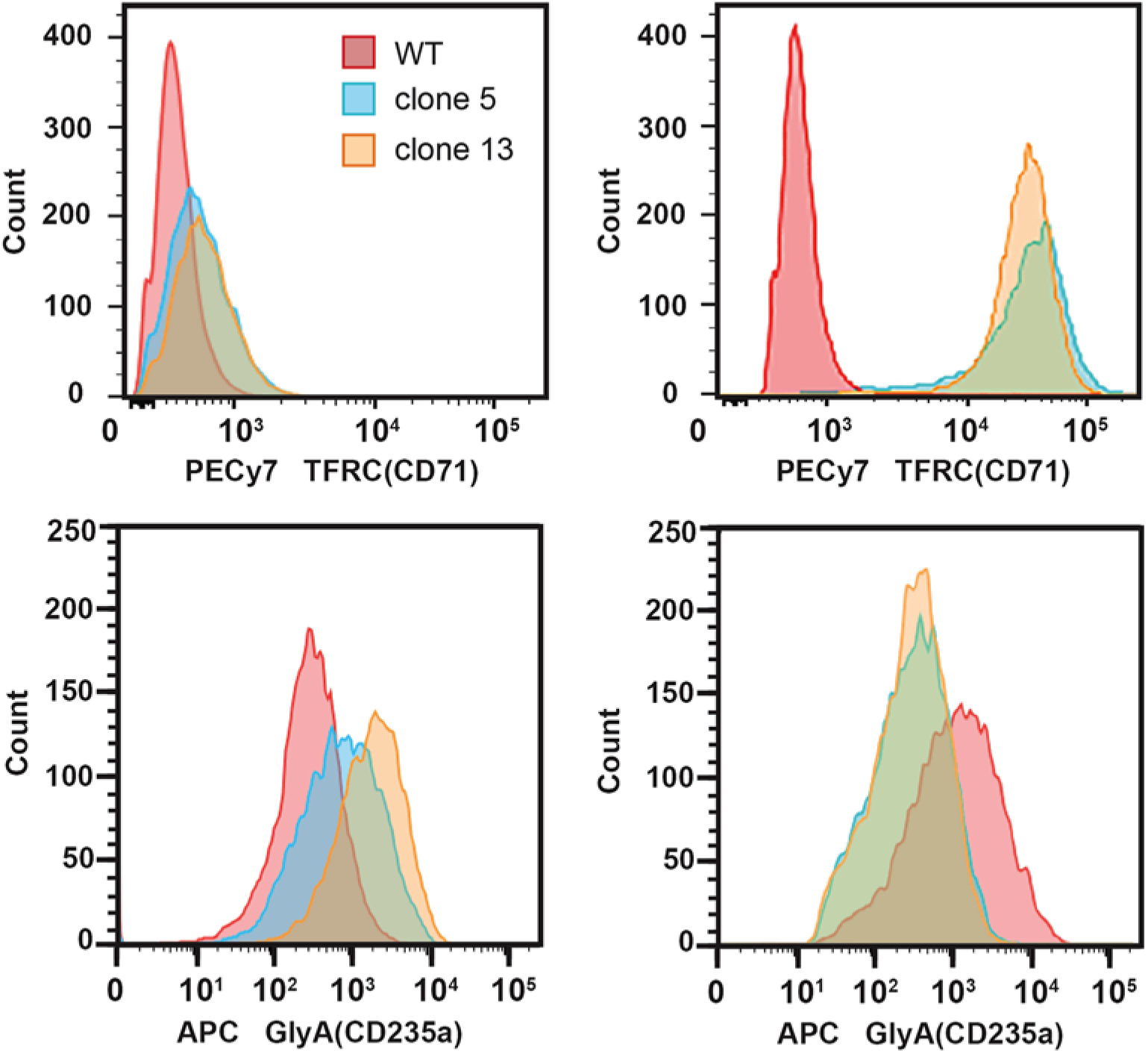
Erythroid maturation is impaired in HEL KO cells compared to WT.

**Table 1.** MnSOD deficiency inhibits erythroid differentiation in response to δALA. Mean fluorescence intensities (MFI) for surface expression of markers of erythroid differentiation, transferrin receptor (CD71) and glycophorin A (CD225a). After 5 days in δALA, wild type HEL cells resemble maturing erythroid cells (low TFRC, high GlyA) whereas *SOD2^−/−^* cells displayed aberrant expression of these biomarkers, consistent with their defect in DNA demethylation which is required for erythroid differentiation. AU – arbitrary units

| (MFI, AU x 10<sup>-2</sup>) | -δALA | +δALA | Fold $\Delta$ |
| --- | --- | --- | --- |
|  | **TFRC** |  |  |
| WT HEL | 2.5 | 5.7 | 2.3 |
| *SOD2*<sup>-/-</sup> cl.5 | 4.4 | 31 | 70 |
| *SOD2*<sup>-/-</sup> cl.13 | 5.0 | 40 | 80 |
|  | **Glycophorin A** |  |  |
| WT HEL | 3.2 | 12 | 3.8 |
| *SOD2*<sup>-/-</sup> cl.5 | 9.0 | 4.5 | -2.0 |
| *SOD2*<sup>-/-</sup> cl.13 | 20 | 4.5 | -4.4 |

## Discussion

We tested the overarching hypothesis that succinate serves as a mitochondrial retrograde signal linking MnSOD activity to nuclear epigenetic reprogramming. Our results suggest a model in which increased mitochondrial O_2_^•-^levels in cells lacking MnSOD leads to inhibition of SDH activity and accumulation of succinate. In turn, this build-up of succinate causes inactivation of the TET family of DNA demethylases. Such disruption of epigenetic homeostasis might contribute to aberrations in developmental pathways or maintenance of cell-type lineage stability. While in this study we have used erythroid differentiation in *SOD2*^−/−^ HEL cells as a phenotypic indicator of disrupted DNA demethylase activity, we also showed disrupted DNA demethylase activity and 5-mC levels in HEK293 cells, thus demonstrating that this effect is not specific to a single cell line or cell type. This data supports the model of an SDH/succinate/TET axis and a role for succinate as a retrograde signaling molecule of mitochondrial origin that significantly perturbs nuclear epigenetic reprogramming to affect cellular phenotype or differentiation state (**Figure 6**). Furthermore, our findings introduce MnSOD as a regulator of the SDH/succinate/TET axis and propose that SDH inactivation in the context of *SOD2* knockout is a result of Fe-S cluster inactivation caused by increased steady-state levels of O_2_^•-^.

**Figure 6.**
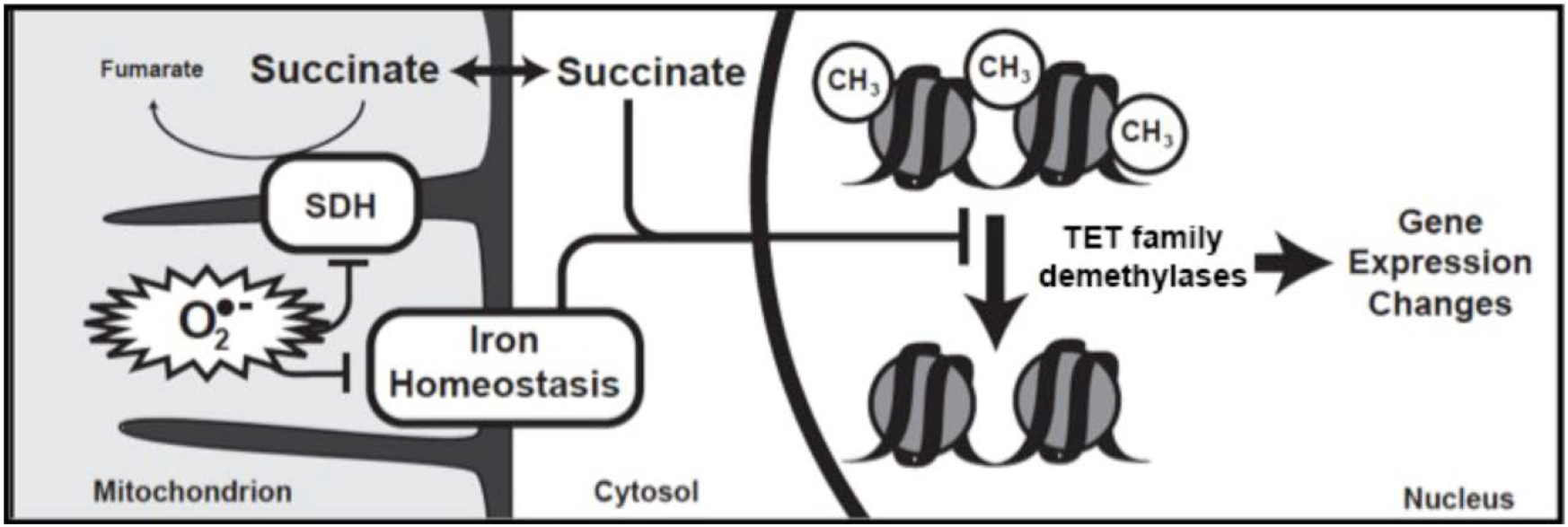
Model depicting a central role for loss of MnSOD leading to higher mitochondrial superoxide which disrupts non-heme iron proteins including SDH resulting in elevated cellular succinate and product-inhibition of α-KG dependent TET enzymes.

Of particular interest will be the continued investigation of the MnSOD/SDH/succinate/TET axis in the context of cancer progression and dedifferentiation. The MnSOD promoter contains a CpG island that is frequently methylated in cancer cells, and MnSOD activity decreases during early carcinogenesis [36–39]. Based on this knowledge, we have proposed an “epigenetic flywheel” in which loss of SOD activity contributes to epigenetic changes, leading to further CpG island hypermethylation of the promoters of *SOD2* and tumor suppressor genes [40]. Findings in this study support the theory that decreased MnSOD activity is linked with increased DNA methylation.

Hypothetically, the epigenetic dysregulation caused by this “flywheel” effect could be initiated at any constituent of the MnSOD/SDH/TET axis. One set of initiators includes SDH mutations. Indeed, SDH mutations, are common in gastrointestinal stromal tumors and paragangliomas and are associated with decreased TET enzyme activity, DNA hypermethylation, poor differentiation, and occurrence of epithelial-to-mesenchymal transition (EMT) [17,41–45]. In addition to genetic mutations, SDH activity can be impaired by cellular iron deficiency induced by various stressors, such as lysosomal dysfunction [46,47].

Relatedly, mutations to fumarate hydratase (FH), another Fe-S cluster-containing protein, or isocitrate dehydrogenase (IDH), lead to accumulation of fumarate and D-2-hydroxyglutarate, respectively, which act in a manner similar to succinate by inhibiting TET and Jumonji histone demethylases, leading to DNA and histone hypermethylation, impairment of homologous recombination [48–50] and failure to differentiate in several contexts [5,51–53]. Interestingly, SDH and IDH mutations alone are sufficient to activate oncogenic programs in the absence of classical kinase mutations by disrupting normal chromosomal topology [54,55].

Another potential set of initiators of the proposed “flywheel” include direct mutations to TET enzymes or differential bioavailability of TET enzyme substrates and cofactors. For example, genetic loss of TET enzymes can lead to increased hematopoietic stem cell self-renewal and eventual development of a myeloproliferative phenotype [56]. Regarding TET substrates, tumor hypoxia has been shown to induce hypermethylation *via* inhibition of TET [57] and O_2_ tension determines cellular differentiation by regulating TET activity in eSCs [58]. Availability of cofactors including Fe^2+^ and ascorbate have also been shown to regulate TET enzyme activity [59–61]. Some of this knowledge is already being translated into cancer therapies. For example, ascorbate treatment has been shown to increase TET enzymatic activity and 5-hmC levels and is associated with reduced tumor progression and/orgrowth in several contexts [62–64]. It should be noted that, while we did not examine the effect of succinate accumulation of histone demethylase activity in this study, we postulate that aberrant histone methylation may also be occurring under these conditions, as KDMs are also α-KG dependent dioxygenases amenable to inhibition by succinate. Such hypotheses will be the focus to future studies.

Ultimately, our results present a link between mitochondrial oxidative metabolism and epigenetic regulation, support the role of succinate in the growing list of metabolic regulators of the epigenome, and introduce MnSOD as a modulator of succinate levels and TET enzyme activity, likely through control of SDH activity.

## Acknwlwdgements

This work was supported by NIH grants R01CA115438, P30CA086862, T32CA078586, and P01CA217797.

## Figure Legends

**Figure 1. Characterization of MnSOD loss in CRISPR/Cas9-edited HEL 92.1.7 cells.** The CRISPR/Cas9 system for creating specific genetic deletions was used to target exon 3 of human *SOD2* in a human erythroleukemia cell line (HEL 92.1.7). Total protein was isolated from clones transfected with the CRISPR/Cas9 expression construct and subjected to Western blot analysis. Two clones were confirmed to show absence of MnSOD protein expression by Western blot analysis.

**Figure 2.***SOD2* **KO cells exhibit decreased activity in enzymes sensitive to inactivation by superoxide and an accumulation of cellular succinate**. **A)** Aconitase activity was measured in a spectrophotometric assay where total cellular protein was combined with NADP^+^ and isocitrate dehydrogenase and the appearance of NADPH was monitored every 30s for 5 minutes. **B)** Succinate dehydrogenase activity visualized via NBT reduction histochemistry was significantly reduced in both 293T and HEL cells without MnSOD expression. **C)** Total cellular succinate increased ~5-fold in both 293T and HEL KO cells. \**p*<.01 compared to WT.

**Figure 3. Activity of TET family of demethylation enzymes is diminished in cells lacking MnSOD.** TET activity was measured in an antibody-mediated colorimetric assay using nuclear protein. A ~2-fold reduction in TET family activity was observed in MnSOD-deficient 293T cells (blue striped bar) and HEL cells (red striped bar). *p*< .05 compared to WT.

**Figure 4. TET enzyme inactivation is associated with perturbed global epigenetic programming. A)** An expected increase in the percentage of genomic 5-methylcytosine was observed in 293T and HEL KO cells. **B)** A corresponding decrease in the percentage of genomic 5-hydroxymethylcytosine was detected in 293T and HEL cells lacking MnSOD expression. \**p*<.01 compared to WT.

**Figure 5. Erythroid maturation is impaired in HEL KO cells compared to WT.** With the addition of δ-ALA, HEL KO cells (clone 5, blue and clone 13, orange) failed to downregulate expression of Transferrin receptor (TFRC) and decreased expression of Glycophorin A (GlyA) compared to WT cells (red) as determined by inspection of AUC.

**Figure 6. Model depicting a central role for loss of MnSOD in disrupting differentiation.** Loss of MnSOD leads to higher mitochondrial superoxide which disrupts mitochondrial non-heme iron proteins including SDH resulting in elevated cellular succinate and product-inhibition of α-KG dependent TET enzymes, resulting in large scale alterations in genomic DNA methylation.

